# Convolutional neural network MRI segmentation for fast and robust optimization of transcranial electrical current stimulation of the human brain

**DOI:** 10.1101/2020.01.29.924985

**Authors:** Carla Sendra-Balcells, Ricardo Salvador, Juan B. Pedro, M C Biagi, Charlène Aubinet, Brad Manor, Aurore Thibaut, Steven Laureys, Karim Lekadir, Giulio Ruffini

## Abstract

The segmentation of structural MRI data is an essential step for deriving geometrical information about brain tissues. One important application is in transcranial electrical stimulation (e.g., tDCS), a non-invasive neuromodulatory technique where head modeling is required to determine the electric field (E-field) generated in the cortex to predict and optimize its effects. Here we propose a deep learning-based model (*StarNEt*) to automatize white matter (WM) and gray matter (GM) segmentation and compare its performance with *FreeSurfer*, an established tool. Since good definition of sulci and gyri in the cortical surface is an important requirement for E-field calculation, *StarNEt* is specifically designed to output masks at a higher resolution than that of the original input T1w-MRI. *StarNEt* uses a residual network as the encoder (ResNet) and a fully convolutional neural network with U-net skip connections as the decoder to segment an MRI slice by slice. Slice vertical location is provided as an extra input. The model was trained on scans from 425 patients in the open-access ADNI+IXI datasets, and using *FreeSurfer* segmentation as ground truth. Model performance was evaluated using the Dice Coefficient (DC) in a separate subset (N=105) of ADNI+IXI and in two extra testing sets not involved in training. In addition, *FreeSurfer* and *StarNEt* were compared to manual segmentations of the MRBrainS18 dataset, also unseen by the model. To study performance in real use cases, first, we created electrical head models derived from the *FreeSurfer* and *StarNEt* segmentations and used them for montage optimization with a common target region using a standard algorithm (*Stimweaver*) and second, we used *StarNEt* to successfully segment the brains of minimally conscious state (MCS) patients having suffered from brain trauma, a scenario where *FreeSurfer* typically fails. Our results indicate that *StarNEt* matches *FreeSurfer* performance on the trained tasks while reducing computation time from several hours to a few seconds, and with the potential to evolve into an effective technique even when patients present large brain abnormalities.

## 1 Introduction

Neurological disorders affect up to one billion people wordwide independently of their age, sex, education or income^1^. In some cases, the quality of life of these people can be substantially improved by a personalized treatment since individual variability has a huge impact on the effectiveness of the therapy. In this study we are going to focus on transcranial electrical stimulation (tES), a neurological therapy that can deliver powerful personalized solutions by stimulating more precisely the target regions or networks of each specific patient.

tES is a non-invasive neuromodulatory therapeutic technique based on the delivery of low electric current via electrodes attached to the scalp that modulate the activity of the brain by exciting or inhibiting target regions. Effects on neuronal excitability depend on the electric field (E-field) distribution [1]. However, and despite recent advances in in-vivo measuring techniques [2], the only approach to predict the E-field in the head with a high spatial resolution is via numerical methods, e.g. the finite element method (FEM) [3]. Such FEM requires geometric information about the tissues in the head, which is obtained from structural images like Magnetic Resonance Imaging (MRI) [4, 5], and relies on knowledge about the electrical conductivity of the different tissues in the brain and the stimulation montage (electrode position, geometry and currents) [6]. This electrical model can then be used to determine the optimal stimulation montage to target specific brain regions and/or networks (Fig. 1) [7, 8, 9, 10]. Therefore, personalized tES requires accurate modeling of the patient’s head to gain knowledge about the E-field induced in the individual brain and predict its effects. The first step in this process is image segmentation, i.e., the classification of each voxel of the image into a specific tissue type of the brain.

**Figure 1:**
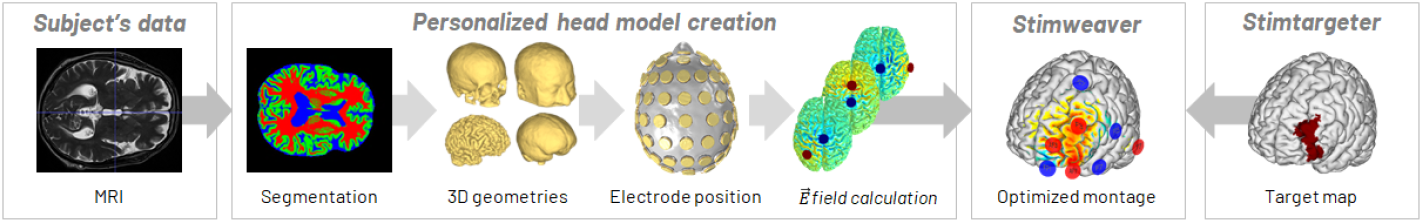
Modeling pipeline used to determine the optimal montage of the electrodes to target a specific brain region: T1-weighted MRI acquisition, segmentation of brain tissues, 3D surface reconstruction, placing of all electrodes, mesh generation, electric field calculation of all possible pairs of electrodes and optimization of the electrode montage based on a target map.

Automated segmentation is a necessary tool for clinical practices due to the fact that manual segmentation has several limitations: it is time consuming and can lead to human errors. The brain has a complex shape, particularly the gray and white matter (GM, WM, respectively), which makes the manual segmentation a difficult task. Nevertheless, automated brain segmentation also faces several challenges. First, a high anatomical and structural variability exists between brains from subjects of different age, sex, ethnic group, and conditions. Second, the segmentation of the WM and GM tissues can often result in masks with low resolution that can lead to “merged” sulci in the final head model, significantly affecting the E-field predictions [5]. Lastly, brain MRI images can have a number of artifacts which can affect image segmentation [11], depending on the scanner and the parameters used to acquire the images.

There are many automated techniques that segment within a few minutes most of the head tissues required for a model (scalp, skull and cerebrospinal fluid (CSF), including ventricles), but GM and WM are more problematic because they are much more complex and computationally demanding for segmentation algorithms. *FreeSurfer* is a standard tool for automated labeling in neuroimaging, specifically for processing and analyzing MRI images of the human brain [12]. Its segmentation pipeline produces WM and GM segmentations that rely on topology corrections and surface deformation based on intensity gradients. *FreeSurfer* results in robust and accurate sulci definition but, on the down side, it requires several hours of computation due to its focus on surface segmentation.

In this work we focus on deep learning methods, and in particular convolutional neural networks (CNN). In the challenging area of brain MRI segmentation, these techniques are being applied to automate the process of brain tissue segmentation, including various brain abnormality segmentation tasks [13, 14]. CNNs have gained a lot of attention in recent years because of their effectiveness in detecting image features not explicitly defined but relevant and highly effective for the medical image segmentation tasks.

### 1.1 Related work

There exist two different types of CNN architectures with their advantages and limitations when focused on our application: voxel-and semantic-wise architectures.

It has been shown that voxel-wise architectures perform worse in cortices consisting of gyri and sulci. This can be explained by the fact that patches centered at boundary pixels contain pixels of multiple tissues. Moreover, these neural networks are very dependent on patch size. Another issue with voxel-wise CNN architectures is the class imbalance between samples or patches since this could indirectly increase the miss-classified patches at the boundaries, which are present in a small amount in the image, or even force an over-training of background samples, which are present in a greater amount. These types of CNN architectures provide a low spatial context, and often use an additional input to improve the performance of the algorithm, such as spatial priors. Training the image segmentation using patches is a slow process; nevertheless, with fewer training images, the model tends to converge faster.

Semantic-wise CNN architectures, on the other hand, train end-to-end and voxel-to-voxel in each slice. Compared to the voxel-wise CNN technique, they generate the entire segmentation from the slice input, which can better use and preserve neighborhood information in the predicted segmentation, and receives as an input the whole image [29, 20, 21, 28]. It is also possible to extract and use patches from the whole image, resulting in the segmentation of the whole patch that is given as input. However, in this case, many discontinuities between patches may appear.

Additional inputs are used to achieve optimal segmentation results, e.g., by incorporating fractional anisotropy (FA), T1 or T2 MRI images in either voxel-wise or semantic-wise architecture.

As can be observed, most of the studies in Table 1 employ a 2D convolution, although some apply satisfactorily a 2.5D convolution or, similarly, a 2D convolution in the sagittal, axial and coronal views. The main limitation of 2D convolution is that it considers the image slice by slice, thus potentially causing discontinuous prediction results across slices. Acquiring 3D contextual information seems to be more natural in 3D brain MRI scans and enables the creation of a deeper and more discriminative neural network. However, this task is faced with memory, data and time limitations. This was addressed in some cases by using small-single 3D patches combined with 2.5D [16] [22] patches as input samples to the neural network or by capturing, only at the initial layer, spatial information with 3D convolutions [21].

**Table 1:**
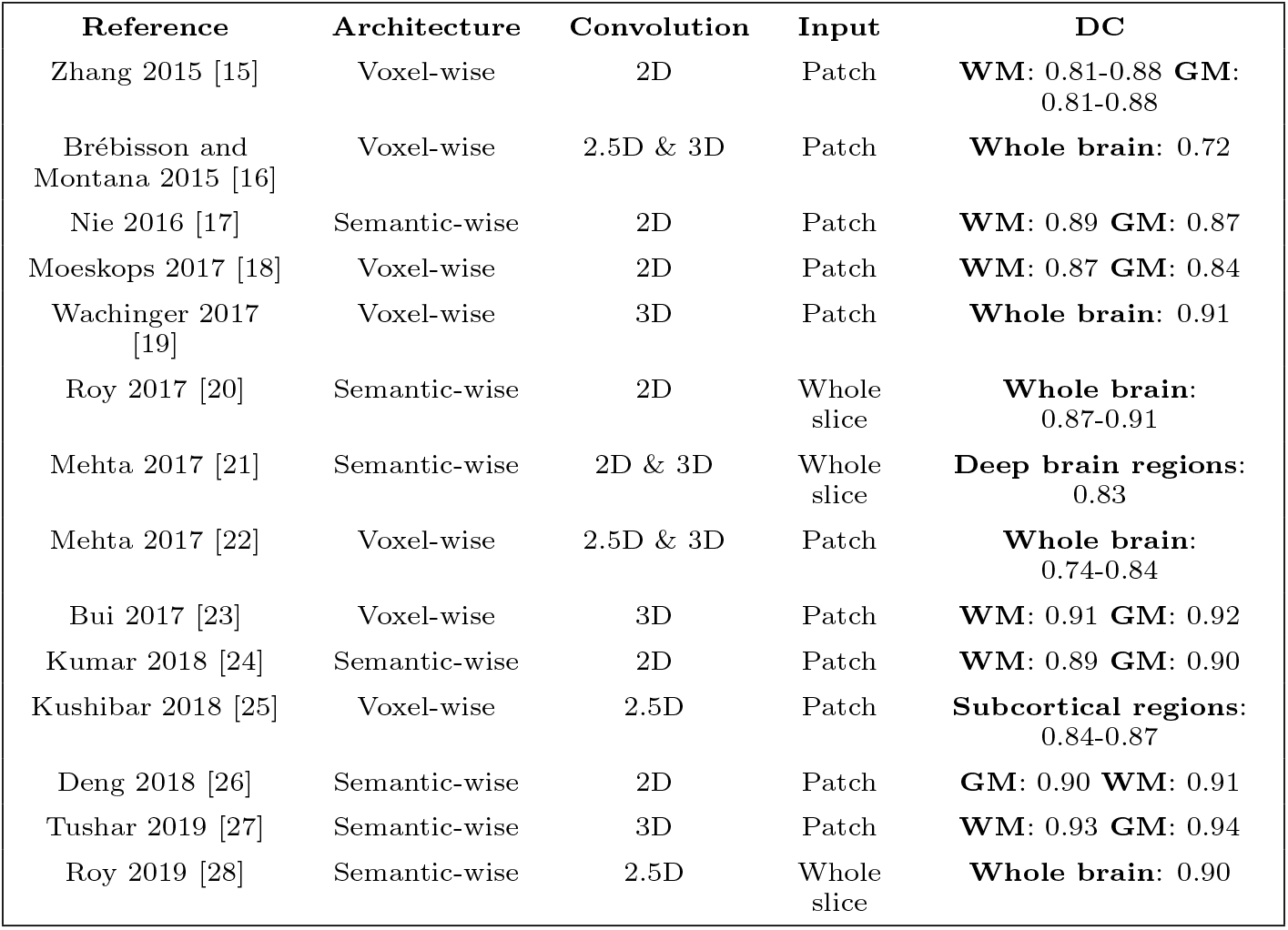
Literature review. Architecture, type of convolution and inputs used in each study, also their performance measured with the Dice Coefficient (DC) metric.

Recently, Leonie Henschel et. al [30] proposed a new *FreeSurfer* pipeline, FastSurfer, based on a full brain segmentation achieved by a semantic-wise CNN to speed-up the process, avoiding the use of skull-stripping and non-linear atlas registration. Although this is an improvement regarding computational cost, still the pipeline requires further postprocessing for topology correction and GM surface creation. In their test a complete *FreeSurfer* run takes approximately 7 hours on the CPU, which can vary depending on image quality, diseases severity... Overall, FastSurfer pipeline achieves the whole brain segmentation in 1 minute on a GPU and the whole surface processing in 5.4 hours on a CPU.

### 1.2 Contributions

In the present work, we present a new technique based on semantic-wise architectures designed to work in a more robust manner that deals with the segmentation of subjects with lesions. The specific requirements for the CNN architecture are:

- Reduce the computation time of the WM and GM segmentations as compared to *FreeSurfer*, but preserving its accuracy.
- Work with just a T1-weighted MRI scan as input, since in many applications a T2-weighted MRI is not available.
- Create WM and GM segmentations with a resolution that has proved to be enough to maintain accurate sulci definition (0.5^3^ mm^3^ resolution), important to generate accurate E-field predictions.

## 2 Methods

### 2.1 Datasets

A sufficient amount of data with representative variability is essential to avoid overfitting and to make the model reliable. We describe next the different datasets used in this work for the training, validation and test of the fully CNN (FCNN) we propose. In Fig. 2 the variability between the datasets used in this work is clearly displayed and in Table 2 further information about the datasets is provided.

**Figure 2:**
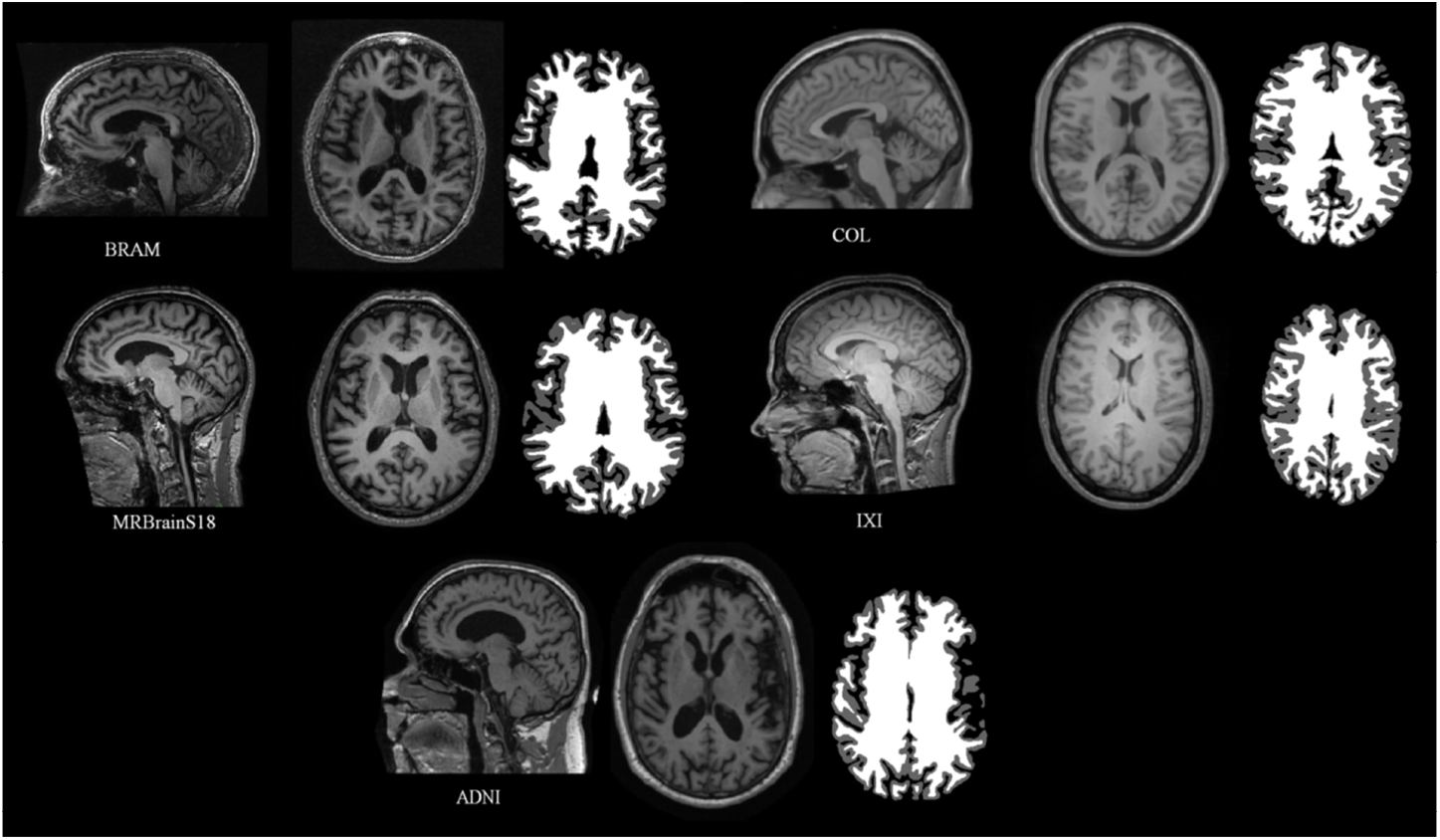
Sample subjects from all the datasets with their corresponding MRI images and volume converted *FreeSurfer* segmentations using iso2mesh [31], highlighting dataset and subject variability.

**Table 2:** Dataset summary. Number of patients, dimensions and resolution of the five datasets called IXI, ADNI, COL, BRAM and MRBrainS18 datasets.

| **IXI** | **T1-weighted MRI scan** | **Segmentations from *FreeSurfer*** | **Manual segmentations** |
| --- | --- | --- | --- |
| Number | 108 | 108 | 0 |
| Dimensions | 256 x 256 x 256 | 512 x 512 x 512 | - |
| Resolution | 1 mm<sup>3</sup> | 0.5<sup>3</sup> mm<sup>3</sup> | - |
| **ADNI** |  |  |  |
| Number | 422 | 422 | 0 |
| Dimensions | 256 x 256 x 256 | 512 x 512 x 512 | - |
| Resolution | 1 mm<sup>3</sup> | 0.5<sup>3</sup> mm<sup>3</sup> | - |
| **COL** |  |  |  |
| Number | 2 | 2 | 0 |
| Dimensions | 256 x 256 x 256 | 512 x 512 x 512 | - |
| Resolution | 1 mm<sup>3</sup> | 0.5<sup>3</sup> mm<sup>3</sup> | - |
| **BRAM** |  |  |  |
| Number | 12 | 12 | 0 |
| Dimensions | 256 x 256 x 256 | 512 x 512 x 512 | - |
| Resolution | 1 mm<sup>3</sup> | 0.5<sup>3</sup> mm<sup>3</sup> | - |
| **MRBrainS18** |  |  |  |
| Number | 7 | 7 | 7 |
| Dimensions | 256 x 256 x 256 | 512 x 512 x 512 | 512 x 512 x 512 |
| Resolution | 1 mm<sup>3</sup> | 0.5<sup>3</sup> mm<sup>3</sup> | 0.5<sup>3</sup> mm<sup>3</sup> |

#### 2.1.1 The IXI+ADNI dataset

This dataset includes 422 3D T1-w MRI scans from the ADNI dataset^2^ with 166×256×256 voxels and size 1.2×0.9×0.9 mm^3^ and also 108 3D T1-w MRI scans from the IXI dataset (see ^3^ for details of scanner parameters) with 256×256×150 voxels and size 0.9×0.9×1.2 mm^3^. Both datasets are normalized to an intensity range of 0 to 255 and converted to 256^3^ dimensions and 1 ^3^ mm^3^ voxel size.

Of the 422 subjects in the ADNI dataset (138 female, mean age 73±4) 234 have mild to severe cognitive impairment and the rest are healthy elderly. Furthermore, this dataset includes MRI images of the IXI dataset, from 3 different hospitals and scanners, and from healthy patients with different sex and ages.

All this data can be used as a T1-weighted MRI input image to train and validate (N=425) the CNN with the aim of optimizing the WM and GM segmentation process, and test (N=105) its performance. The desired output data (ground truth) is obtained from *FreeSurfer* using the 3D T1-weighted MRI scan. The surface of the brain tissues given by *FreeSurfer* is converted to volume using iso2mesh [31], a MATLAB toolbox. The volume is created with higher resolution and dimensions, 0.5^3^ mm^3^ and 512^3^ respectively. This resolution, as discussed previously, is crucial when creating the mesh and modeling the E-field.

#### 2.1.2 The COL and BRAM datasets

We will in addition use datasets not used during training, namely COL and BRAM, for out-of-sample testing.

COL is an average of 27 T1-weighted MRI scans of the same individual (age around 28). The two MRI scans (1 female) have 160×211×195 dimensions and 1^3^ mm^3^ resolution. The BRAM dataset (4 males) was collected for a previous study in a pilot, double-blinded, block-randomized, sham-controlled trial of tDCS intervention (NCT…) [32] in older adults with slow-gait and mild-to-moderate executive dysfunction. Written informed consent was obtained and all study procedures, and all recruitment material, were approved by the Hebrew SeniorLife Institutional Review Board. Some of these images were used for this work and had 256×256×162 dimensions and 0.9×0.9×1 mm^3^.

All MRI scans and *FreeSurfer* segmentations are processed exactly as the IXI+ADNI dataset.

#### 2.1.3 The MRBrainS18 dataset

In order to compare the performance of *FreeSurfer* and our proposed DCNN model, the MRBrainS18 dataset with manual segmentations is also used in this study. T1-weighted MRI scans (256×256×192 voxels and 0.9×0.9×3.0 mm^3^ resolution) of the MRBrainS18 dataset (see ^4^ for details of scanner parameters) and their corresponding manual segmentations (240×240×48 voxels and 0.9×0.9×3.0 mm^3^ resolution) were acquired from 7 patients with diabetes, dementia and Alzheimers, and others with increased cardiovascular risk with varying degrees of atrophy and white matter lesions (age > 50).

Again all MRI scans and *FreeSurfer* and manual segmentations are processed exactly as the other datasets.

### 2.2 Preprocessing

To level out the dependency of CNN performance from data quality, a preprocessing pipeline is designed (Fig. 3):

1. Bias field correction to correct inhomogeneities in the magnetic field of the MRI machine.
2. Global contrast normalization to normalize the contrast between 0 and 1.
3. Registration to align each image with an asymmetric T1-weighted MRI template of 0.5^3^ mm^3^ resolution and 394×466×378 dimensions, built from the average of a number of unbiased nonlinear images in the MNI152 database^5^. The alignment is implemented using an affine transformation. The final dataset has also the dimensions and resolution of the template independently from their original values.
4. Cropping to remove background and reduce the dimensions of our image to 394×394×339.
5. Histogram matching using as a template the average of the intensity values in each image pixel of only the training dataset.

**Figure 3:**
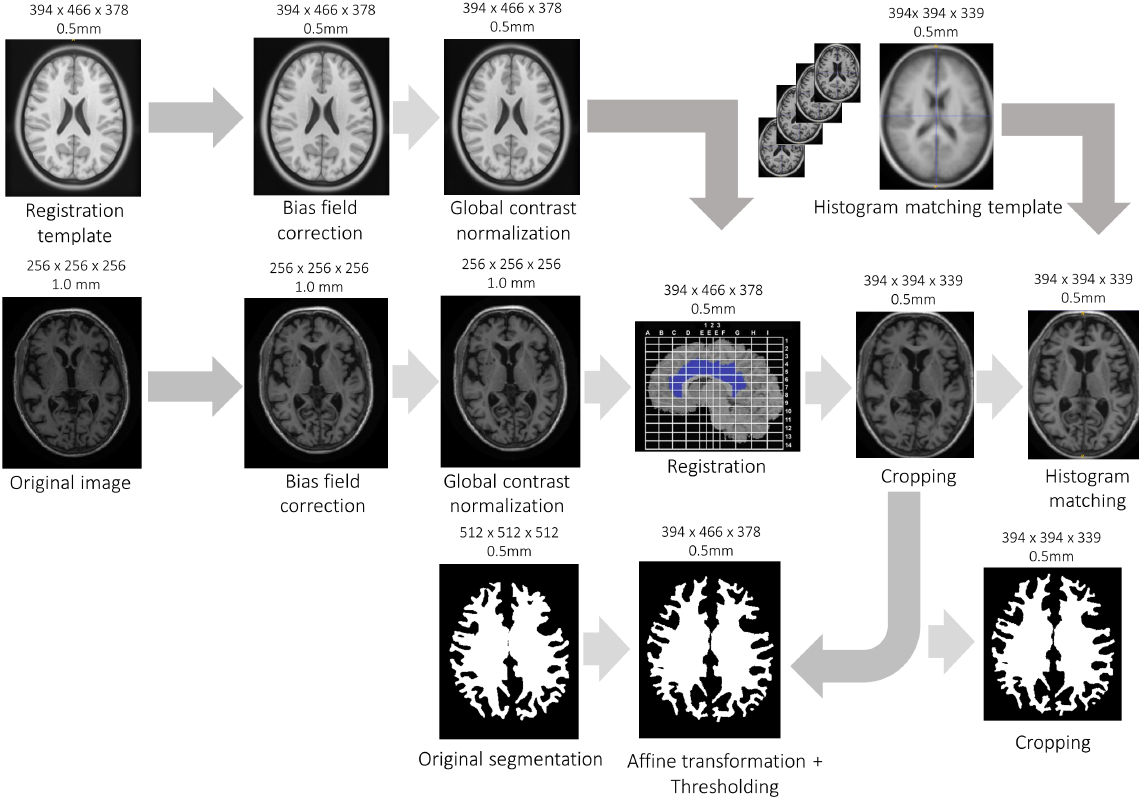
Preprocessing pipeline of one subject 3D MRI implemented with Python programming language. **a)** Pre-processing of the registration template, **b)** average of the training dataset to get the histogram matching template and **c)** preprocessing of the MRI to be segmented. Dimensions and resolution of the data in each stage are indicated.

The ANTs^6^ tool (Advanced Normalization Tools for image processing), is used for bias field correction, registration and histogram matching. Each subject requires about 4 minutes to be preprocessed, with the registration step being the most time consuming. The WM segmentation used as ground truth includes the ventricles, which are later removed.

### 2.3 Network architecture

The chosen architecture (Fig. 4) is a U-net [33] with a ResNet [34] encoder.

**Figure 4:**
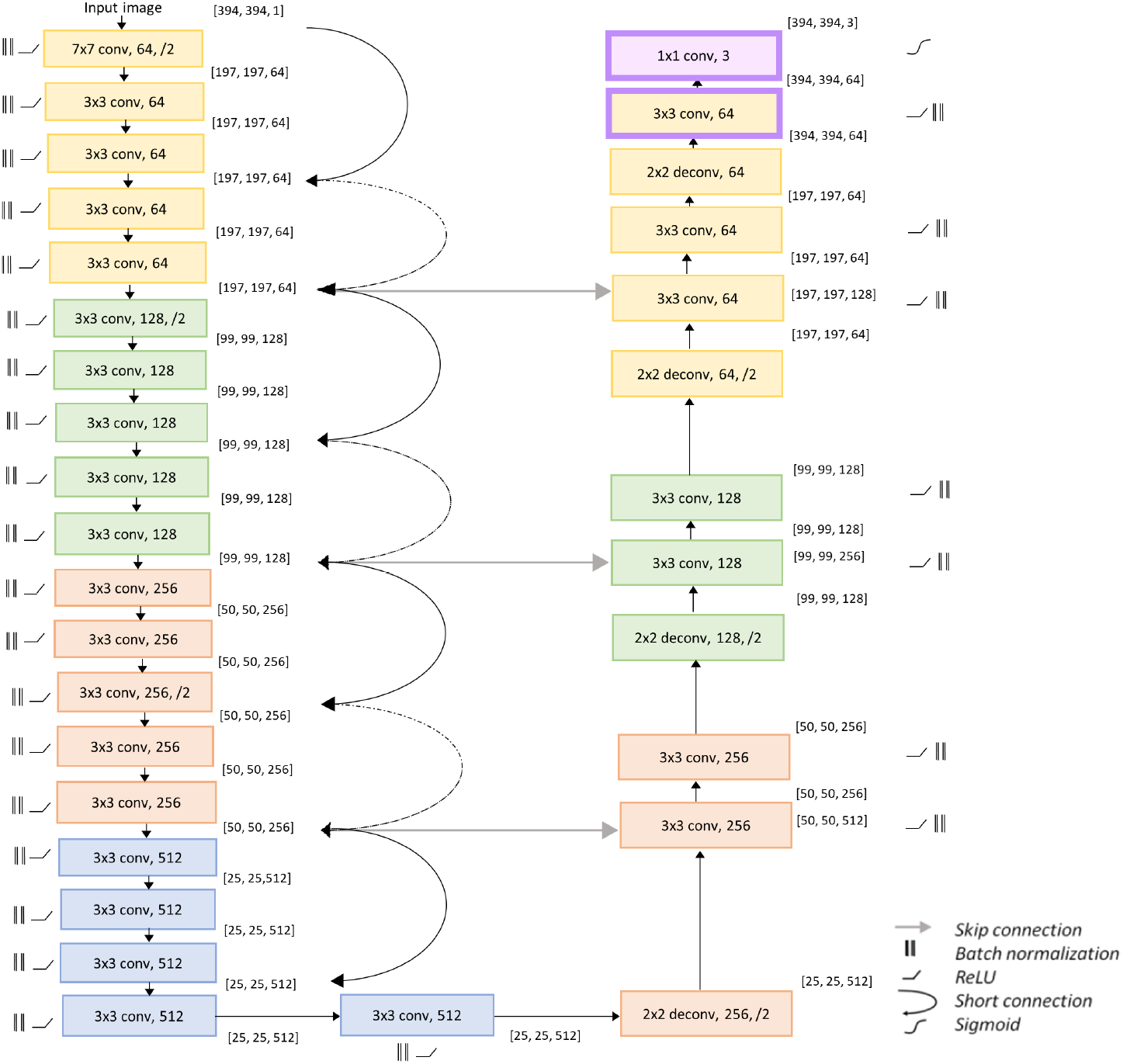
*StarNEt* architecture, using *ResNet* (in our implementation it had 34 layers) as the encoder and U-Net skip connections. The two last layers of the decoder (in purple) represent the case in which a prior is considered.

The encoder of our model corresponds to the *ResNet-34* architecture pre-trained with *ImageNet*. In *StarNEt*, the encoder applies 2D convolutions with 3×3 kernels and stride 1 in several layers. Each of these convolutions is followed by batch normalization and the non-linearity of a rectified linear unit (ReLU) activation function. The encoder is divided into four blocks with a different channel dimension. When the number of features is increased, the same 2D convolution is applied but with a stride of 2, reducing the dimensions of the convolution output and preserving the spatial information, information that would be lost if using a pooling layer. The particular property of *ResNet* is the presence of short connections between layers, which sum the values of previous layers to the current one to avoid the “vanishing gradient” problem. Linear projection by the shortcut connections can be performed to match the dimensions when needed.

The decoder of the architecture deconvolves the inputs received from the encoder with a 2 x 2 kernel size and a stride of 2. Then, the output of the deconvolution receives the features detected by the encoder in the different blocks with the skip connections from *U-net*. The same convolutions used in the convolutional pathway are used in the deconvolutional pathway. The final output of the decoder is convolved with a 1 x 1 kernel, followed by a sigmoid activation function. Afterward, an output with the same dimensions as the initial input is obtained, but with the corresponding background, WM, and GM segmentations.

#### 2.3.1 Fine-tuning

*StarNEt* implements fine-tuning as a transfer learning strategy to retrain the weights of the encoder, by continuing the backpropagation with the new dataset acquired for our study. The encoder, trained earlier with the *ImageNet* dataset, is available online^7^. However, the decoder is initialized with random weights.

#### 2.3.2 Information about slice location

Since contextual information about the third (vertical or axial dimension) is missing, the numeric identifier of the 2D slice is provided as input to be taken into account. An additional channel with the slice number normalized (from 0 to 1) is input to the last two layers before the convolution is applied, to lead to the *StarNEt* model with prior (Fig. 4).

### 2.4 Loss function

The loss function chosen to be optimized is cross-entropy^8^, also called logarithmic loss function. Cross-entropy (CE) penalizes the false negative values and false positive values, but especially those predictions that are confident and wrong. It is defined by

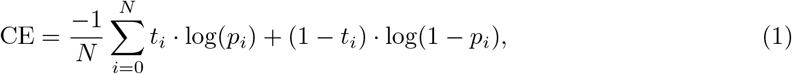

where *t_i_* is the true label, *p_i_* the predicted label for pixel i and N corresponds to the total number of pixels of each input x.

### 2.5 Evaluation metric

In this work, the segmentation accuracy is measured with the DC, which calculates the overlap ratio between the automatic segmentation and ground truth. The DC value evaluates the segmentation accuracy and integrates measures of over and underestimation into a single metric. It is estimated by

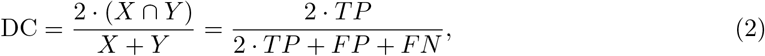

where TP, FP, and FN are true positives, false positives and false negatives respectively. X values are the predictions and Y values are the true labels.

### 2.6 Implementation and environment

PyTorch^9^ is the open-source machine learning library for Python used for the implementation of the model learning with the IXI+ADNI dataset (Fig. 5). The stochastic gradient descent (SGD) optimization is performed with Adam [35] and The batch size of 10 slices is constrained by the 11 GB of memory of the NVIDIA GeForce RTX 2080 Ti GPU. The learning rate is kept to 1·10^−3^ during the whole training. Finally, the model with the highest validation metric is chosen and re-trained for 12 epochs with a lower learning rate (3·10^−5^) to finish the convergence.

**Figure 5:**
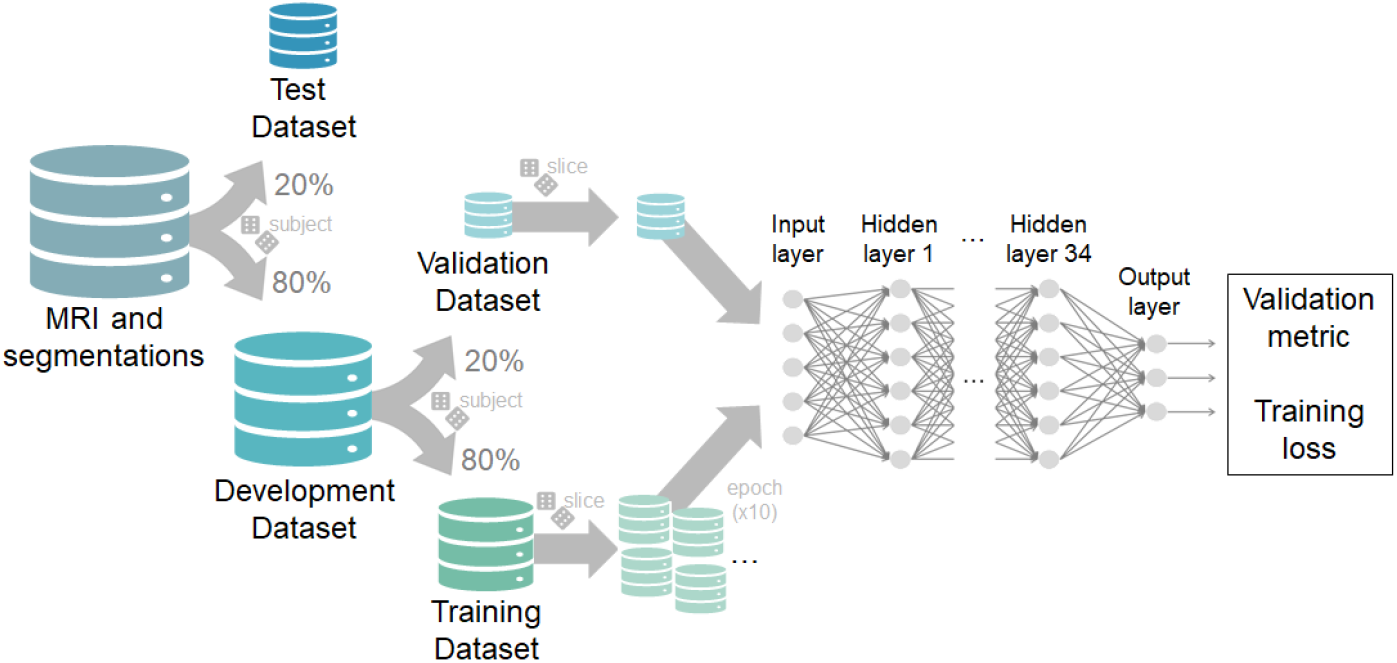
Model training workflow. The data is randomly split into the test and development dataset (20% and 80% respectively) by subjects. Then, the development dataset is divided in the same way into two main groups: the validation and the training dataset (20% and 80% respectively). The training set is shuffled randomly by slice and is given to the model in batches of 10 slices. This process is repeated 10 times (10 epochs). In each epoch the validation set evaluates the performance of the model.

### 2.7 Electrode montage optimization

In order to test the impact of our approach on electrode montage optimization two head models are built using the T1-weighted MRI of the female subject from the COL dataset. In both models the images are segmented into CSF (the ventricles were included by manually segmenting with ITK-SNAP ^10^ [36]), air filled sinuses, skull and scalp using SPM 8 ^11^ with the MARS package [37] and also into WM and GM (including cerebellar WM and GM) but using *FreeSurfer* in one model and *StarNEt* in the other. Then, the segmentations are combined using MATLAB ver. R2018b so as to make sure that the tissues are surrounded at least by 1mm layer of another tissue, to generate smoothed surfaces of all tissues and resolved any intersection between them. All of this is performed using MATLAB’s Image Processing Toolbox and iso2mesh [31].

Later, an optimization algorithm (*Stimweaver* [8]) is applied to target a common cortical area in many tES applications: the left dorsolateral prefrontal cortex (lDLPFC) [38]. The algorithm considers the *E_n_* (normal E-field of the cortex), as this component is thought to predict excitability changes of pyramidal cells. It requires a predefined matrix of 64 bipolar montages with a common reference electrode that can be linearly combined to generate the E-field induced by any montage using these electrodes.

The 64 electrodes are placed on the scalp on positions defined by the 10/10 EEG system [39]. The PISTIM Ag/AgCl ^12^ electrodes (1 cm radius, 3.14 *cm*^2^ area) with conductive gel underneath are modeled as cylinders.

After setting the appropriate tissue conductivities for DC-low frequency range: 0.33 S/m, 0.008 S/m, 1.79 S/m, 0.40 S/m, 0.15 S/m and 10^−5^ S/m respectively for the scalp, skull, CSF, GM, WM and air [5]. The model is solved in Comsol 5.3, the E-field normal to the GM surface (*E_n_*) is calculated for all the bipolar montages with Cz as a cathode (−1 mA) and each of the other electrodes as the anode (+1 mA).

The optimization is constrained for the maximum current per electrode (2.0 mA) and total injected current (the sum of all the positive currents: 4.0 mA). A genetic algorithm is employed to find the solution with a maximum of 8 electrodes.

The metrics used to fit the optimization problem are the Weighted Cross-Correlation Coefficient (WCC) and the Error Relative to No Intervention (ERNI) (mV^2^/mm^2^) [8], which sums over cortical mesh nodes:

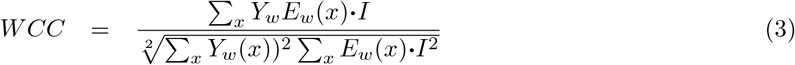

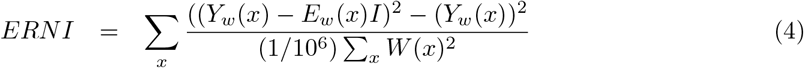

*I* is the array of electrode currents, *Y_w_* (*x*) and *E_w_*(*x*) correspond respectively to the desired and the output electric field for each mesh point, and *W*(*x*) is the weight set to each node dependent on the local region.

While WCC measures the related weighted correlation coefficient of the target map and the E-field, a number between −1 and 1, ERNI takes into account the overall quality of the solution over the cortical surface. In order to obtain the relative ERNI (RERNI), the value of the ERNI is scaled with its optimal solution (when *Y_w_*(*x*))^2^ — *E_w_*(*x*) = 0):

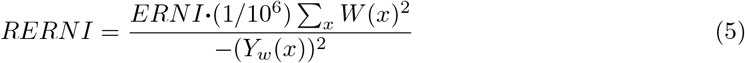

Then the results of the optimizations were compared in terms of electrode positions, currents and average E-field on target.

### 2.8 Study with MCS patients

*FreeSurfer* does not successfully segment brain tissues in some patients with brain trauma, such as in minimally conscious state (MCS). Due to some brain abnormalities the voxels of the WM and the GM are misclassified. Here *StarNEt* with prior is tested for segmenting these brain tissues in such cases.

A dataset with T1-w-MRIs of 18 MCS patients (10 females, mean age 34±15, 5 with traumatic brain injuries (TBI), 6 with TBI and skull defects, 7 with neither TBI nor skull defects), was provided by the Coma Science Group of Université de Liegè. The dataset was acquired using a 3 Tesla scanner (Siemens Tim Trio; Siemens Medical Solutions, Erlangen, Germany) with 120 T1-weighted slices (repetition time [TR] = 2300 ms, echo time [TE] = 2.47 ms, flip angle 9 degrees, voxel size = 1×1×1.2 mm^3^, field of view [FOV] = 256 mm^2^). These scans are also processed using our preprocessing pipeline (section 2.2).

The data was used in accordance with local Research Ethics Committee and all patients’ next of kin or legal representative confirmed their approval and consent to participate in the study and the reuse of these fully anonymized data for investigation proposes.

## 3 Experiments and results

### 3.1 Training and validation with the IXI and ADNI datasets

The evolution of the first and second training for *StarNEt* with and without prior can be seen in Fig. 6 and Fig. 7.

**Figure 6:**
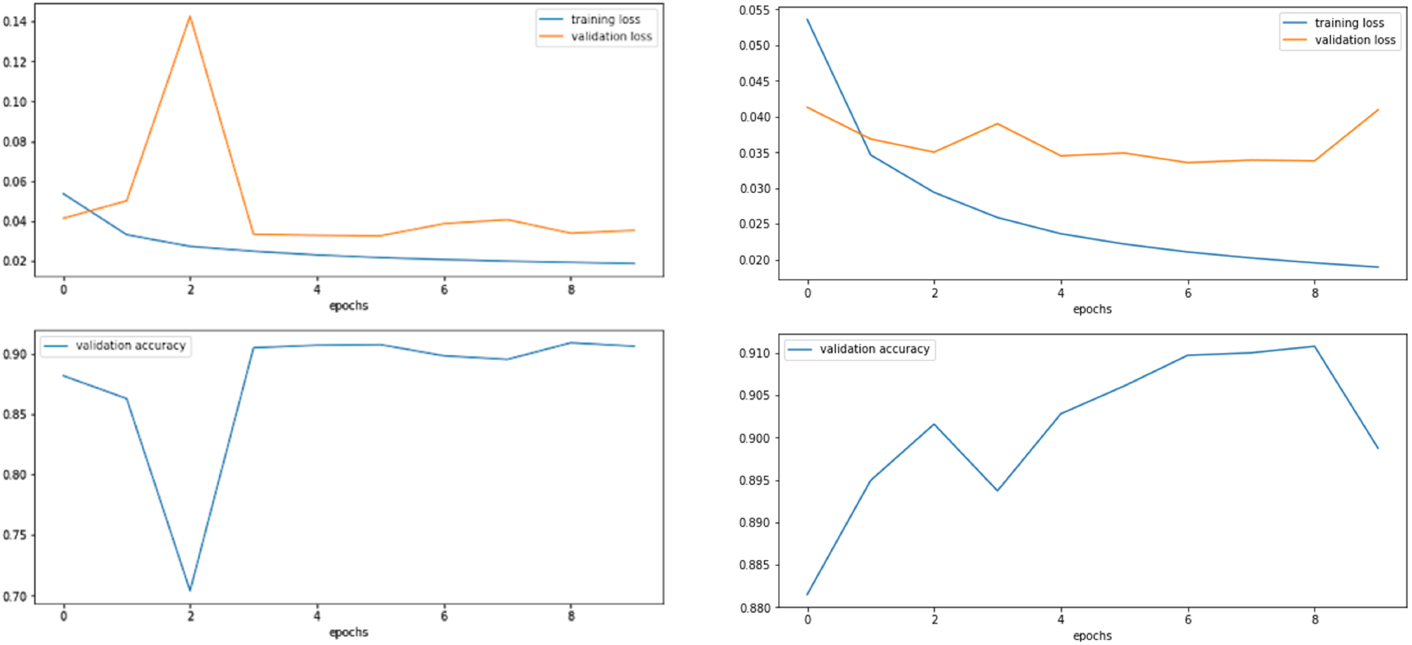
First training evolution without (left) and with prior (right) with a learning rate of 1·10^−3^. Loss value of the training dataset and loss value of the validation dataset in each epoch of the training. The validation accuracy, the DC value, is also represented.

**Figure 7:**
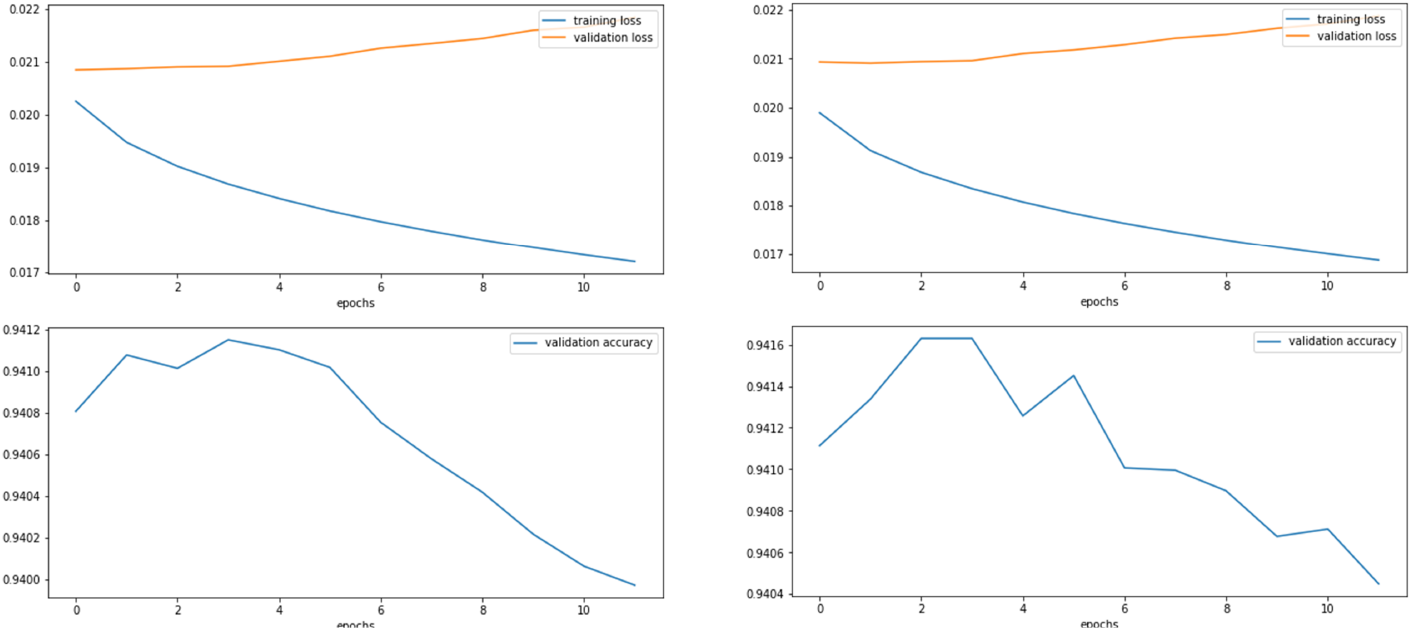
First training evolution without (left) and with prior (right) with a learning rate of 3·10^−5^. Loss value of the training dataset and loss value of the validation dataset in each epoch of the training. The validation accuracy, the DC value, is also represented.

The DC of the WM and GM segmentations for the model without the slice identifier that gives the best validation metric value is 0.957 and 0.933 respectively (0.945 in average). The model with the 3D prior results in a DC of 0.958 (WM) and 0.935 (GM) with an average of 0.947. It can be seen that the slice number does not make a great impact on the final result of the validation dataset and both have in common that it is harder to segment GM pixels than WM ones.

### 3.2 Test with the IXI, ADNI, COL and BRAM datasets

Table 3 demonstrates that, with a DC around 0.89, *StarNEt* performs better on the IXI+ADNI dataset than on the out-of-sample datasets (COL and BRAM). However, the performance with the COL and BRAM datasets does not decrease too much.

**Table 3:** Performance of the model with and without slice number information using the validation IXI and ADNI datasets. The DC of both WM and GM segmentations are annotated in each case.

| StarNet with no prior |  |  |  |  |
| --- | --- | --- | --- | --- |
| Training data | Metric | Test data |  |  |
| IXI and ADNI datasets | DC | IXI and ADNI datasets | COL dataset | BRAM dataset |
|  | WM | 0.922 | 0.891 | 0.882 |
|  | GM | 0.862 | 0.828 | 0.793 |
|  | Average | **0.892** | **0.860** | **0.837** |
| StarNet with prior |  |  |  |  |
| Training data | Metric | Test data |  |  |
| IXI and ADNI datasets | DC | IXI and ADNI datasets | COL dataset | BRAM dataset |
|  | WM | 0.919 | 0.915 | 0.883 |
|  | GM | 0.854 | 0.826 | 0.789 |
|  | Average | **0.886** | **0.870** | **0.836** |

Again, GM segmentation is less accurate than WM segmentation. This difference increases with the out-of-sample COL and BRAM datasets and is confirmed by the 3D reconstructed segmentation in Fig. A1. In Fig. A1, discontinuities across slices were more visible in the GM-CSF interface. The simple prior studied in this work does not help the neural network to learn data not used in the training, with the exception of the COL dataset, in which an improvement is observed.

By providing a view of several regions of the brain, Fig. 8 shows that *StarNEt* is less accurate at the boundaries of the WM and GM.

**Figure 8:**
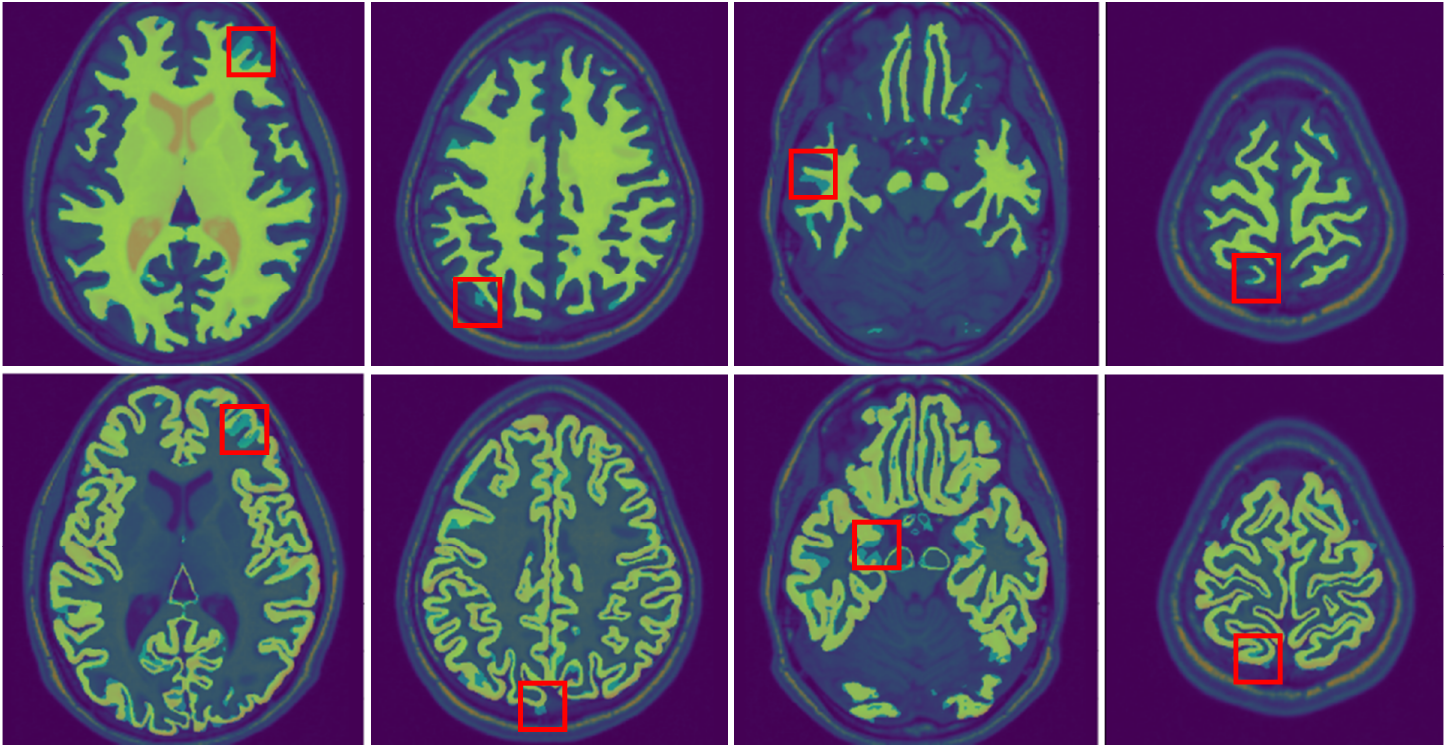
Segmentation of a few WM (top) and GM (bottom) slices from a COL female subject superposed over the ground truth, which is generated by *FreeSurfer*. True positive pixels are colored in light green and false positive and false negatives in light blue (some highlighted in red). Behind there is the corresponding MRI.

In Fig. 9 there are some examples where *FreeSurfer*, used as ground truth, gives sometimes inaccurate segmentations.

**Figure 9:**
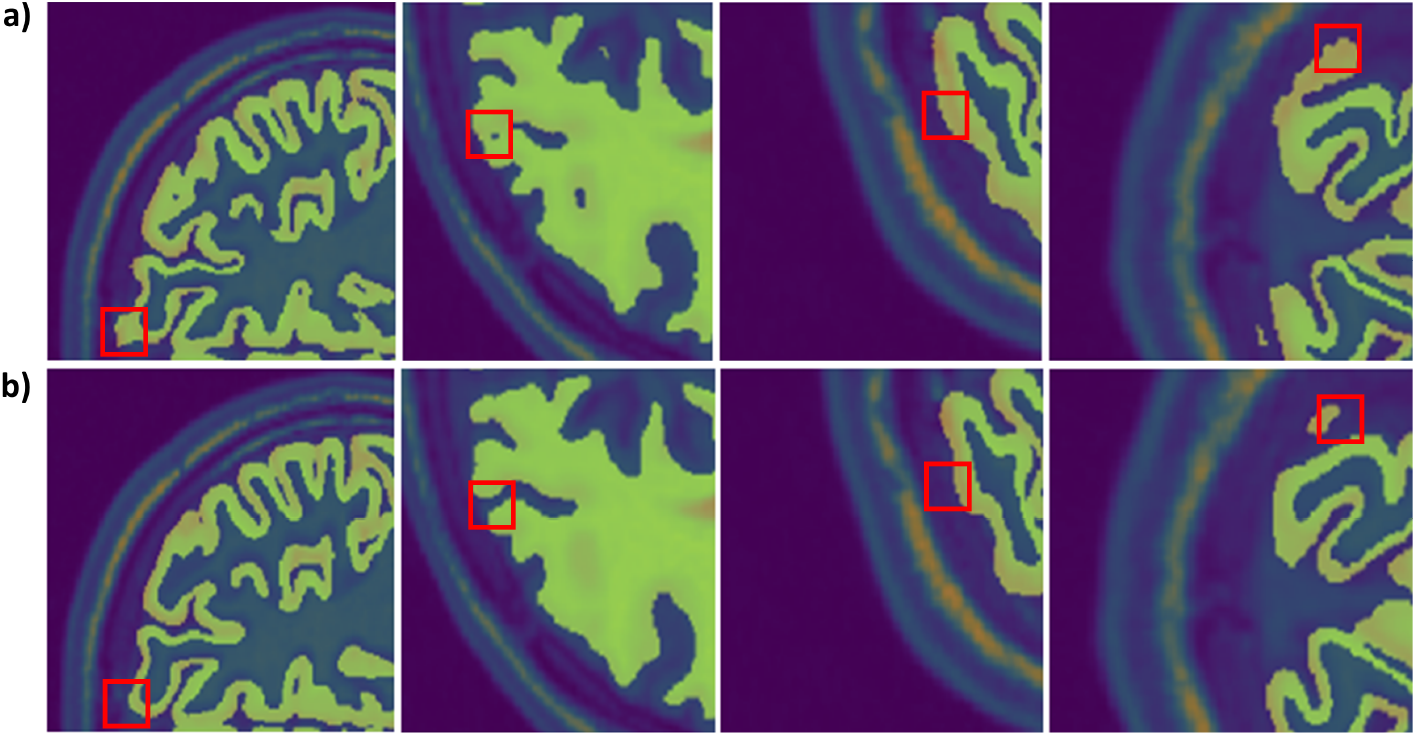
Regions where **a)** *FreeSurfer* performance is overcome by *StarNEt* **b)** with no prior.

### 3.3 Test with the MRBrainS18 dataset

In this section, the performance of *FreeSurfer* on a slice basis is compared with the model that considers the slice number and the one that does not, by using manual segmentations as ground truth. The computation time to segment each of the subjects in MRBrainS18 dataset using *StarNEt* was 4 seconds and several hours using *FreeSurfer*, both having a comparable accuracy, with a DC average of around 0.7.

In Table 4 the performance of both *FreeSurfer* and *StarNEt* is presented in more detail. *FreeSurfer* is significantly less accurate on GM segmentations, which may also explain the reduced accuracy for *StarNEt* in GM pixel classification with respect to WM. In addition, *StarNEt* is slightly more accurate segmenting GM than *FreeSurfer*.

**Table 4:** WM and GM DC of *StarNEt* (with and without prior) and *FreeSurfer* for the MRBrainS18 dataset using manual segmentations as ground truth.

| DC | *FreeSurfer* | *StarNet* with<br>no prior | *StarNet* with<br>prior |
| --- | --- | --- | --- |
| WM | $0.740 \pm 0.025$ | $0.731 \pm 0.046$ | $0.744 \pm 0.026$ |
| GM | $0.659 \pm 0.046$ | $0.668 \pm 0.038$ | $0.669 \pm 0.040$ |

By comparing the outcomes obtained from *StarNEt* with prior and without prior slice number information (Table 4), we see that slice location information enables the model to classify better the WM, allowing *StarNEt* to beat *FreeSurfer* on both tasks (WM and GM segmentations).

Results in Fig. B1 manifest that, in general, the slices of the 3D MRI scan that are in the middle surpass in performance the first or last slices in both *FreeSurfer* and *StarNEt*. These slices can be mostly improved by *StarNEt* with prior, however as these are a few, they do not influence the final DC value much (Table 4).

### 3.4 Montage optimization derived from *StarNEt* and *FreeSurfer* segmentation head models

In Figure 10, the electrode montages for both optimizations is shown, proving that the distribution of the normal component of the electric field (*E_n_*) created by the two montages, which are also nearly the same, is very similar. We have the same scenario in terms of total and maximum current for electrode injected (see Table 5).

**Figure 10:**
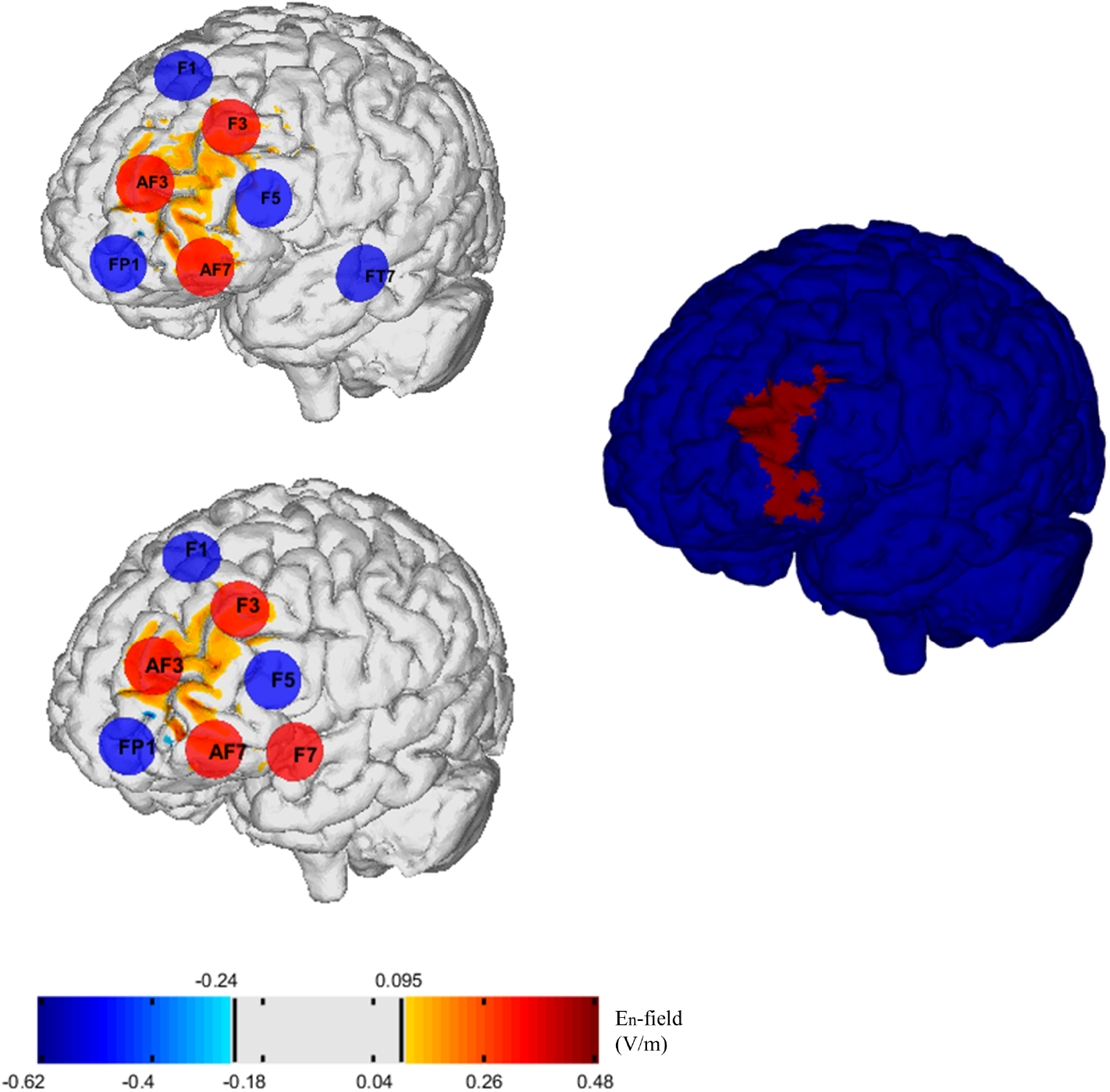
*E_n_* field induced by optimized montages using *FreeSurfer* (top) and *StarNEt* (bottom) WM and GM segmentation (bottom). The optimization target was the lDLPFC, as shown in the right panel.

**Table 5:** Total current (Total I), maximum current (Max. I), the WCC and the relative ERNI (RERNI) for the standard head model using FreeSurfer segmentations and for the case using *StarNEt* segmentations.

|  | Standard | *StarNet* |
| --- | --- | --- |
| Total I (mA) | 3997 | 3998 |
| Max. I (mA) | 1697 | 1795 |
| WCC | 0.51 | 0.55 |
| RERNI | 0.26 | 0.30 |

As seen in Table 5 the WCC values obtained in both cases are really close to each other (7.84% increase) and the same for the relative ERNI (15.38% increase), which demonstrates that both head models reached a good optimal solution.

The results are also analyzed in more detail in terms of average E-field (*E_n_*) in Table 6 for the closest regions to the target (weight=10), the lDLPFC, and for the furthest ones (weight=2). The E-field generated for both montages in their corresponding model are similar between each other, proved also before in Fig. 10. Then, if we try each montage for the opposite head model we observe that in this case the Standard model has a higher average E-field than using its Standard montage for local target regions. Oppositely, the *StarNEt* model shows lower E-field values using the Standard montage.

**Table 6:** Average E-field in the normal component (*E_n_*) in local (w=10) and non-local target regions (w=2) for the two montages and in the two models created using *FreeSurfer* or *StarNEt* WM and GM segmentations.

|  | Standard montage | *StarNet* montage |
| --- | --- | --- |
| Standard model | w=10 0.062 V/m<br>w=2 -0.001 V/m | w=10 0.079 V/m<br>w=2 -0.001 V/m |
| *StarNet* model | w=10 0.053 V/m<br>w=2 0.000 V/m | w=10 0.068 V/m<br>w=2 0.000 V/m |

### 3.5 *StarNEt* applied to images from MCS patients

Since we did not have manual segmentations from these patients as groundtruth we evaluated the results obtained from *StarNEt* with visual inspection. We could see a clear improvement (Fig. 11) in some of them when compared to *FreeSurfer* performance by only visual quality evaluation, suggesting that *StarNEt* is more robust segmenting data from subjects with lesions. Specifically, the improvement was more relevant with patients that had more damage, in both TBI and skull defects and TBI without skull defects groups. In the first, half of the patients lead to better results using *StarNEt* and 3/5 in the second. Finally, with patients without TBI and without skull defects *Star-NEt* gave similar results except for the two subjects shown in Fig. 11.

**Figure 11:**
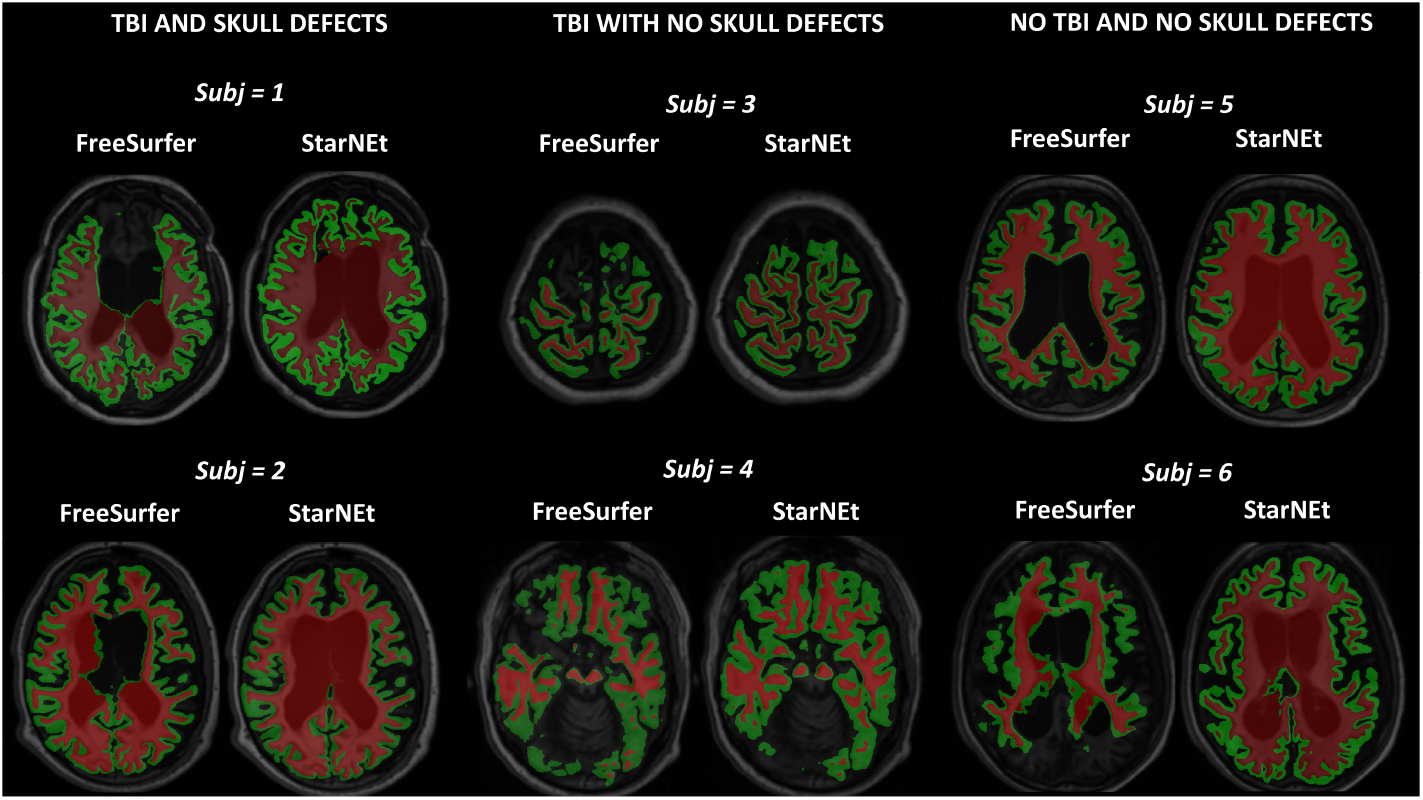
Sample images from 6 of the WM and GM segmentations obtained from each of the MCS patients using *FreeSurfer* or *StarNEt*. Only one slice from the 3D MRI is shown. The data from the first column belongs to the group of TBI and skull defects, the data from the second column belongs to the group of TBI with no skull defects and the last one to neither TBI nor skull defects group.

## 4 Discussion

The benefit of using semantic-wise architectures is that they provide good performance since contextual spatial information is better preserved as opposed to voxel-wise architectures. Furthermore, voxel-wise architectures, in spite of making algorithm slower, involve choosing a patch size, adding complexity to the problem. Semantic-wise architectures do have one drawback: they need a greater amount of training data. This is not a serious limitation in our work (we have availability to a large number of images).

*StarNEt*, a semantic-wise architecture using only a T1-weighted MRI scans, implements 2D convolutions slice by slice, which requires less data, time and memory than 3D convolutions, and achieves a DC of 0.92 for the WM and 0.85 for the GM with the IXI+ADNI dataset.

To our knowledge, Roy et al. [20] created the first architecture that, by combining *U-Net* and *SegNet*, segmented the whole brain slice by slice and opted for simply applying 2D convolutions. Error corrective boosting was the main key to achieve good performance but, apart from this, they did not need any additional input to get the desired results. The advantage of *StarNEt* with respect to their model is that it uses a pretrained neural network available online, which makes the neural network training converge rapidly. Apart from that, it relies on the slice number in order to consider the third dimension lost with 2D convolutions. This prior gives encouraging results but can probably be fine-tuned (the implementation approach was simple).

*StarNEt* outperforms the results reported in the literature (see Table 1) with the exception of Tushar et al. [27], who obtained the best results by using a semantic-wise architecture as well. They also relied on the *ResNet* architecture, but the difference is that they used 3D convolutions to patch inputs, a fact that slows down the process. Bui et al. [23] also achieved better results than *StarNEt*, but their voxel-wise architecture uses multi-modal images. All the studies that beat *StarNEt* have in common the use of 3D convolutions, including Wachinger et al. [19], who also used multi-task learning to compensate for the lack of spatial information of voxel-wise architectures. Some resulted in a better GM segmentation [24] [26], but this might be due to the use of *FreeSurfer* segmentations as ground truth which, as discussed before in results, yields worse performance on GM segmentations than WM ones. Moreover, the disadvantage of the first one [24], is that they implemented an architecture that requires multi-resolution inputs to adequately function. Finally, Roy et al. [28] also obtained marginally better results than *StarNEt* using 2.5D convolutions.

Despite the good results obtained with *StarNEt*, it should be noted that the dataset used in the other studies was different and the number and type of tissues classified were also different. Apart from that, the ground truth used to train the model in this work is obtained from *FreeSurfer* and is not manual segmentation. This is a limitation, since the DC could penalize some pixels that were classified well by *StarNEt* just because *FreeSurfer* ground truth was inaccurate there.

Another important result is that, by applying previously the preprocessing pipeline designed in this work, the model performs properly with datasets not seen during training—the BRAM and COL datasets. This last test of the model in a separate dataset is not usually performed in other studies and it is crucial to see the generalization ability of the model.

There is no doubt that *FreeSurfer* is less efficient than *StarNEt* in computational time. *StarNEt* can provide the segmentation of the whole MRI volume of one patient in a few seconds while *FreeSurfer* takes several hours.

To be consistent with the affirmation that *StarNEt* performs similarly to *FreeSurfer*, we decided to compare their performance using manual segmentations from MRBrainS18 as ground truth. *FreeSurfer* produced a DC of 0.740 and 0.659, while *StarNEt* resulted in a DC of 0.744 and 0.670 for the WM and GM respectively. The small improvement was possibly due to the prior that considers the slice number of the input since *StarNEt* with no slice prior was less accurate. As seen before in the results section, the slices that benefit from this prior are only few, which might be the reason for which the total DC does not change greatly. However, it is possible that this slice number identifier would have been more helpful and beneficial for the performance of the model in the presence of less data to train *StarNEt*.

This work was motivated by the need for a fast segmentation technique with the same performance as *FreeSurfer*. Although image preprocessing takes around 4 minutes, the proposed technique does not require hours and instead segments the scans in 4 seconds on average in the test machine (regular, high line PC with graphics card). *StarNEt* works with just a T1-weighted MRI scan of the patient and, independently from its resolution and dimensions, generates a segmentation of 0.5^3^ mm^3^ resolution, essential to model the effect of the electric field of tES in the brain model. Although the segmentations are obtained slice by slice with *StarNEt* and consequently some discontinuities across slices were found, the entire 3D WM and GM segmentations have been reconstructed and found to have quite good sulcus definition. This was confirmed by the results of the optimizations with the two head models which were also similar both in terms of electrode positions, currents and average E-field on target. The *StarNEt* model shows lower E-field values using the Standard montage and we hypothesize that this fact might be due to the deeper sulcus in the model created by *FreeSurfer* segmentations (Standard model), their architecture/conductivity, notably CSF, may create a higher induced E-field [4].

Segmentation of brain MRI data from patients with brain defects following trauma or surgery is sometimes challenging, since *FreeSurfer* can fail to segment WM and GM tissues properly as proved before in results. *StarNEt* has shown to give improved segmentation results for some of the worst cases.

## 5 Conclusions

Here we presented StarNEt, a 2D semantic-wise architecture that, by using only a T1-weighted MRI scan, has a strong generalization ability to segment WM and GM human brain tissues. Furthermore, it is significantly faster than FreeSurfer, reducing the computational time from hours to seconds (not including the 4 minutes of preprocessing), and can provide better segmentations for both WM and GM, when using the slice number as 3D prior. StarNEt has also proved to be a practical tool for tDCS head modeling, since it can output segmentations masks with 0.5^3^ mm^3^ voxel size (essential for accurate prediction of the E-field) from T1-weighted MRI of any resolution. TDCS electrode montage optimization using WM and GM segmentations from *StarNEt* gives, in fact, comparable results to segmentations from *FreeSurfer*. Finally, *StarNEt* is especially valuable for segmentation in presence of GM or WM large lesions.

## 6 Other Potential Directions

- The first potential future direction for *StarNEt* is for the segmentation of additional tissues and structures in the brain. This refers to the segmentation of not only the WM and GM, but also the CSF, skull and scalp, which should be easier to segment than WM/GM due to their simpler shape. In fact, the segmentation would be more natural, given that now the ventricles could be considered as ventricles, and not as WM, and the other tissues could be considered as brain tissues and not background. Thus each specific structure could be classified by a different label and performance expected to improve. Finally, the segmentation of lesions could be also considered.
- In this project, a model with 2D convolutions has been studied, but an evolution to true 3D convolutions and segmentation should be considered as well in order to overcome WM and GM discontinuities between slices after the 3D reconstruction. One of the main limitations to be faced is GPU memory (many more parameters to optimize) and data availability. To address memory limitation, multiple GPU could be used to segment the entire MRI image with volumetric convolutions. Finally, instead of changing to a 3D convolution, one could consider the possibility of using also the neighbor slices of the slice that wants to be segmented as inputs for *StarNEt*.
- Re-training *StarNEt* with manually segmented data would make the model more accurate, especially for classifying GM voxels, segmented worse by *FreeSurfer* than WM voxels. Since they are more difficult to acquire we could also use dataset augmentation to increase the training dataset.
- In order to further personalize the neural network to different types of patients, one could investigate a method to transfer information to the neural network about sex, age or neurological condition of the subject, in a manner similar to what was performed with slice number.
- Finally, another line of work would be to compare the proposed model with existing ones using the same datasets, which would provide better benchmarking.

## Acknowledgments

This project has received funding from the European FET Open project Luminous (European Union’s Horizon 2020 research and innovation programme under grant agreement No 686764). The results and conclusions in this article present the authors’ own views and do not reflect those of the EU Commission. We acknowledge Alzheimer’s Disease Neuroimaging Initiative (ADNI) (National Institutes of Health Grant U01 AG024904) and DOD ADNI (Department of Defense award number W81XWH-12-2-0012) for providing part of the data collection, https://brain-development.org/ for sharing their data for this project and specially, we would like to acknowledge all members of the Coma Science Group for the data acquisition as well as the Radiology department of the University of Liege.

---

**Figure A1:**
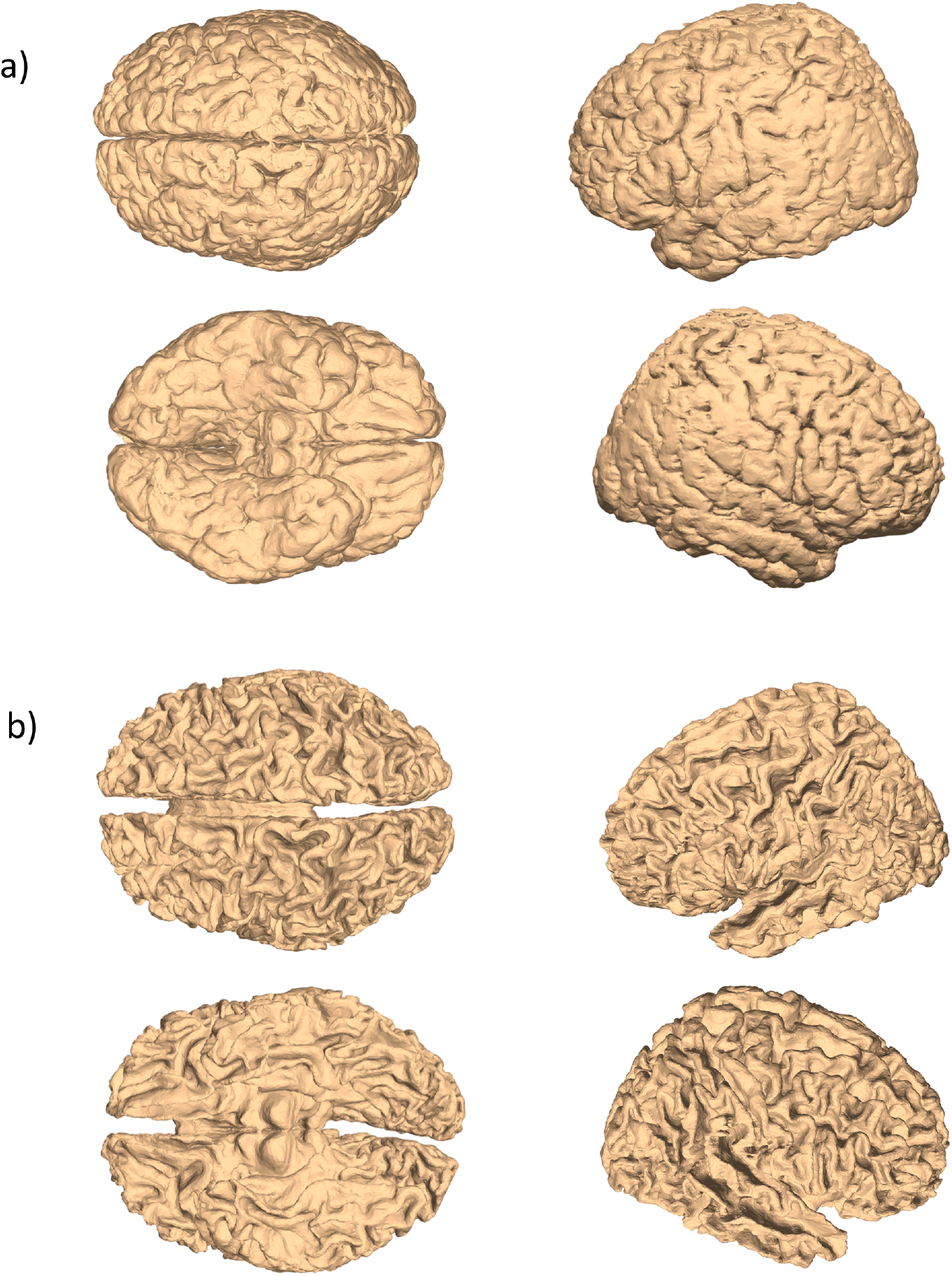
GM (top) and WM (bottom) surface meshes (0.5^3^ mm^3^ resolution) of COL male in 4 different views. They are obtained from the segmentations obtained with StarNEt with prior.

**Figure B1:**
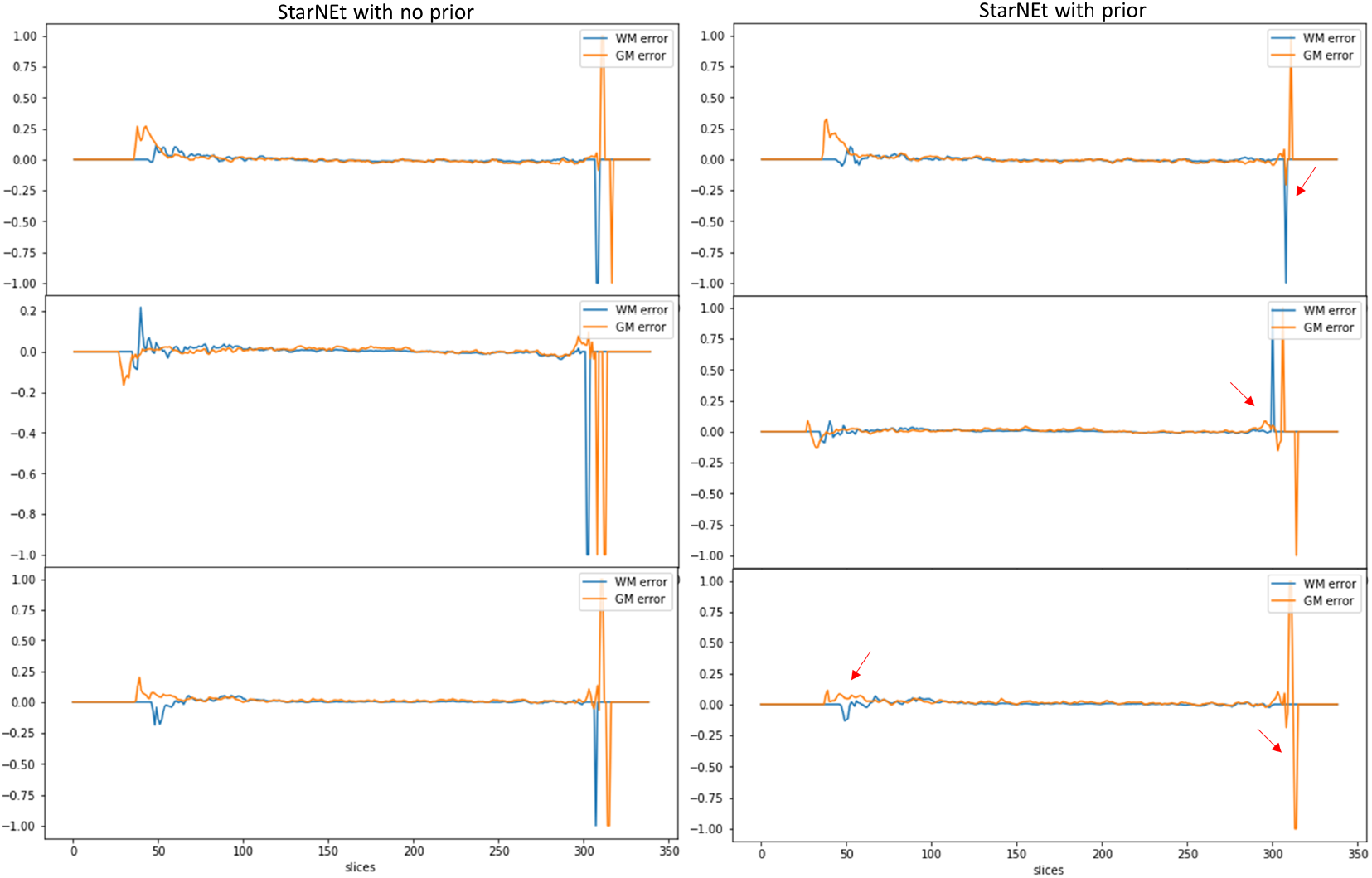
DC error between FreeSurfer and StarNEt for each slice in 3 subjects (top-bottom) from the MRBrainS18 dataset using StarNEt with and without prior for the WM and the GM case: whether the error value is > 0 means that FreeSurfer is worse on the task than StarNEt and vice versa. The more visible improvement or worsening indicators of StarNEt with prior are specifically marked with a red arrow in the plot.

## Footnotes

1 https://www.who.int/mediacentre/news/releases/2007/pr04/en/

2 http://adni.loni.usc.edu/

3 http://brain-development.org/ixi-dataset/

4 https://mrbrains18.isi.uu.nl/data/

5 http://www.bic.mni.mcgill.ca/ServicesAtlases/ICBM152NLin2009

6 https://sourceforge.net/ps/advants/

7 https://pytorch.org/docs/stable/_modules/torchvision/models/resnet.html#resnet34

8 https://ml-cheatsheet.readthedocs.io/en/latest/loss_functions.html

9 https://pytorch.org/

10 http://www.itksnap.org

11 (https://www.fil.ion.ucl.ac.uk/spm/software/spm8/

12 https://www.neuroelectrics.com/products/electrodes/pistim/

